# Predicting Protein-Ligand Binding Structure Using E(n) Equivariant Graph Neural Networks

**DOI:** 10.1101/2023.08.06.552202

**Authors:** Ashwin Dhakal, Rajan Gyawali, Jianlin Cheng

## Abstract

Drug design is a costly and time-consuming process, often taking more than 12 years and costing up to billions of dollars. The COVID-19 pandemic has signified the urgent need for accelerated drug development. The initial stage of drug design involves the identification of ligands that exhibit a strong affinity for specific binding sites on protein targets (receptors), along with the determination of their precise binding conformation (3-dimensional (3D) structure). However, accurately determining the 3D conformation of a ligand binding with its target remains challenging due to the limited capability of exploring the huge chemical and protein structure space. To address this challenge, we propose a new E(n) Equivariant Graph Neural Network (EGNN) method for predicting the 3D binding structures of ligands and proteins. By treating proteins and ligands as graphs, the method extracts residue/atom-level node and edge features and utilizes physicochemical and geometrical properties of proteins and ligands to predict their binding structures. The results demonstrate the promising potential of EGNN for predicting ligand-protein binding poses.

## I. Introduction and Background

The pursuit of new drugs involves a substantial investment of time, resources, and funding. The lengthy process of drug discovery, spanning approximately 12 years and costing around 2.6 billion dollars per drug [1], showcases the challenges inherent in this field. When the outbreak of COVID-19 occurred, the world faced a race against time to find effective treatments and vaccines. The devastating consequences of the pandemic highlighted the critical importance of expediting the computational-based drug development process to save lives and prevent the chaos caused by such viral outbreaks [2], [3]. Furthermore, the vast number of possible drug-like molecules, estimated to be on the order of 10^60^ [4], far exceeds experimental capabilities, necessitating the utilization of computational methods to narrow down the search space.

Accurate prediction of protein-ligand binding structures is fundamental to structure-based drug design. By understanding the interactions between a protein and a ligand, researchers can identify lead compounds with high binding affinity and potential therapeutic efficacy (refer to **Figure 1** to comprehend the docking of a ligand to its target receptor). This knowledge allows for the rational design of new drugs and facilitates the optimization of their properties to improve treatment outcomes.

**Figure 1:**
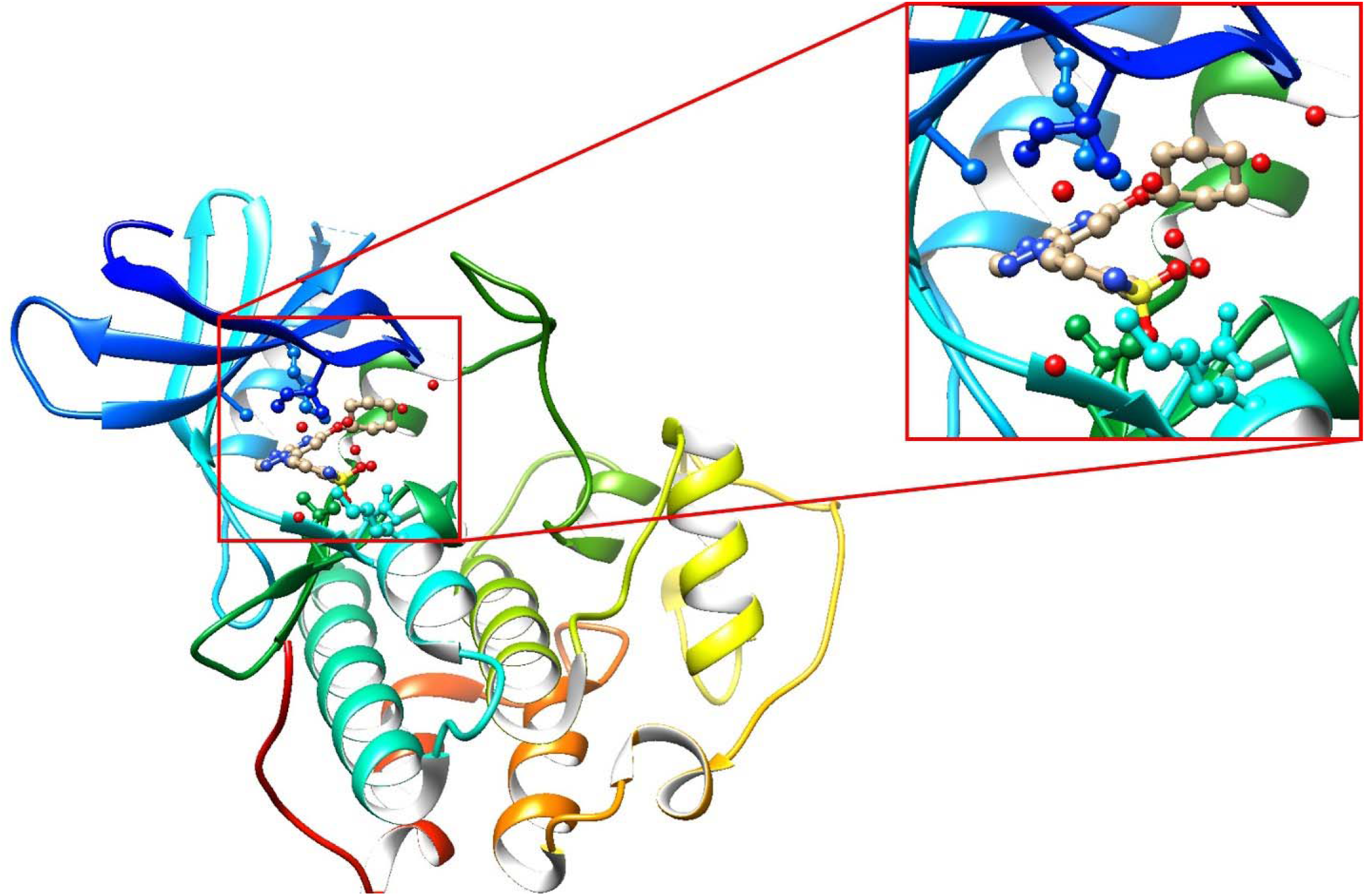
Structure of the protein-ligand complex formed by the docking of triazolopyrimidine inhibitor to the single chain Human Cyclin-dependent kinase 2 (PDB ID: 2c6m [7]). Protein shown in rainbow color and ligand shown in ball and stick representation.

Small molecule ligands play a crucial role in modulating the biological functions of proteins [1], [5]. Consequently, accurate prediction of protein-ligand binding structures is fundamental in the field of computational drug design. Molecular docking, a widely used method, aims to predict the position, orientation, and conformation of a ligand when bound to a target protein, thus providing insights into the potential therapeutic effects of the ligand [6]. However, the inherent flexibility of protein side chains and the impact of solvents pose challenges to traditional docking procedures, necessitating the development of improved methods.

One of the major challenges in protein-ligand binding prediction lies in the inherent flexibility of proteins and the conformational changes that binding pockets undergo to accommodate diverse ligand scaffolds [8]. Protein side chains exhibit dynamic motion, and the role of solvents further adds complexity to the binding pocket geometry. Conventional molecular docking procedures often fail to account for these conformational fluctuations, limiting their prediction accuracy [5]. Addressing this challenge is crucial to improve the reliability of binding pose predictions.

The computational expense associated with exploring the vast search space of protein-ligand interactions is a significant hurdle in drug design. Traditional methods rely on exhaustive search algorithms that can be computationally demanding and time-consuming [1]. Given the staggering number of possible drug-like molecules and the need to assess their interactions with thousands of protein types, there is a pressing need for computationally efficient approaches to accelerate the screening process. Deep learning-based methods have shown promise in reducing the search space, but their accuracy and speed remain areas of active research.

Computational approaches, including deep learning techniques, offer the potential to significantly reduce the molecular search space in drug discovery. However, to be truly effective, these methods must be both accurate and computationally efficient to scan the vast biological and chemical spaces for desired and unexpected effects. With thousands of protein types in the human proteome, computational screening of potential interactions is essential before conducting in vitro and in vivo testing. The ability to rapidly and accurately predict protein-ligand binding structures is crucial for structure-based drug design and the estimation of binding free energy [9].

Over the years, several computational methods have been developed to address the challenges of protein-ligand docking. Traditional docking methods primarily focus on calculating binding energy or affinity, followed by ranking the ligands based on their scores [9]. However, manual verification is often necessary due to the limitations of these methods. Recent advancements in computational drug design have introduced deep learning-based approaches that offer faster runtimes compared to traditional search-based methods. Nevertheless, improvements in accuracy remain a key objective [1].

The complex nature of protein flexibility, the computational expense of exploring the search space, and the significance of binding pose determination necessitate the development of advanced computational methods.

This research introduces a novel approach for predicting the optimized binding pose of protein-ligand interactions using an EGNN [10]. Our method incorporates both atomic and spatial features of proteins and ligands to improve the accuracy of pose prediction. By leveraging the power of EGNN, we aim to overcome the limitations of existing methods and provide a more comprehensive and effective solution for computational drug design.

## II. Literature review

Precise prediction of how proteins and ligands interact has long been a significant challenge in the field of structure-based drug design. While techniques like cryo-electron microscopy (Cryo-EM) [11] [12] can provide detailed structural information about protein-ligand complexes, they are expensive and not always readily accessible for all cases. As a computational alternative, molecular docking has emerged as a powerful tool with significant implications for drug design, including the discovery of potential hits and leads, as well as subsequent optimization efforts. These approaches rely on scoring functions and optimization algorithms to assess the correctness of proposed structures or poses. However, due to the vast search space and rugged landscape of scoring functions, these methods tend to be slow and inaccurate, particularly for high-throughput workflows [1].

Molecular docking faces challenges, leading to increased usage of AI/ML methods for protein-ligand interactions. In 2015, Ashtawy and Mahapatra introduced a machine learning approach to estimate the difference between predicted and true protein-ligand complex structures [13]. Their prediction aimed to select the optimal pose of a ligand within a protein’s binding site, utilizing physicochemical and geometrical properties of protein-ligand interactions. They found that machine learning models trained to predict RMSD (Root-Mean-Square Deviation) values outperformed standard scoring systems.

Similarly, Grudinin et al. [14] tested a regression-based method for scoring binding poses in the 2015 D3R Challenge. They trained their model using regularized regression and affinity and structural data from the PDBBind database [15]. More recently, a strategy based on reinforcement learning was presented by a team led by Jose for estimating pose scores. In this approach, the agent learns not only to identify the binding site but also to optimize the optimal position for maximum efficiency. The network for reinforcement learning-based protein-ligand docking includes a GraphCNN layer that incorporates atomic/molecular properties as a feature vector.

The introduction of EQUIBIND [16] in 2022, inspired by their previous geometric & graph deep learning based model for protein-protein interaction: EQUIDOCK [17], has brought about a significant paradigm shift in the field. This approach utilizes SE(3)-Equivariant Graph Neural Networks to predict drug binding poses. EQUIBIND’s model exhibits excellent empirical performance compared to the most recent state-of-the-art baselines. Other recent works by Stark et al. and Lu et al. have also developed deep learning models that treat docking as a regression problem, predicting the binding pose in a single shot. Although these methods are significantly faster than traditional search-based approaches, they have yet to demonstrate substantial improvements in accuracy.

TANKBind [18], developed by Lu et al., has made progress by independently predicting docking poses for each potential pocket and ranking them using an interatomic distance matrix. However, the performance of these one-shot or few-shot regression-based methods still falls short of traditional search-based methods.

In the pursuit of molecular docking advancements, DIFFDOCK [19], a diffusion generative model (DGM), has been proposed. This model defines a diffusion process over the degrees of freedom involved in docking, including the position of the ligand relative to the protein, the ligand’s orientation within the binding pocket, and the torsion angles describing its conformation.

The development and evolution of these various approaches have shed light on the challenges associated with protein-ligand binding pose prediction. The goal of this project is to create an Equivariant Graph Neural Network for predicting protein-ligand interactions (PLI). This approach has the potential to greatly reduce the need for resources, time, and costs associated with experimental physical tests when screening for new treatments.

## III. Methodology

### 1. Data Acquisition and Processing

In this study, we employed the publicly available PDBBind dataset [15], a valuable resource containing a comprehensive assemblage of experimentally determined binding affinity data. Specifically, we utilized the most recent version of the dataset, PDBBind-v2020, which encompasses binding information, including K_d_, K_i_, and IC_50_ values, for a substantial collection of 19,443 distinct protein-ligand complexes. From this extensive pool, we selected a subset of 1,000 protein-ligand complexes to form the basis of our experiment.

To facilitate the subsequent analysis, we first split the prepared PDBBind-v2020 dataset into three distinct sets: a training set, a validation set, and a test set. The training set was constructed by randomly assigning 750 protein-ligand complexes, while the validation set and test set were composed of 200 and 50 complexes, respectively.

Once the data was partitioned, we proceeded with the crucial step of data preprocessing. Given that our Graph neural network necessitated both the true structure of the complex and a randomly oriented structure as input, we employed Pymol [20], a widely-used software tool, for this purpose. Importantly, we utilized Pymol’s non-GUI interface, which allowed us to flexibly manipulate the orientations of the ligands and proteins. During this preprocessing stage, we implemented random translation and rotation operations on the ligands and proteins along the positive and negative axes of the three-dimensional Cartesian coordinate system (x, y, and z). Specifically, we prescribed a range of random rotation angles spanning from 1 to 360 degrees, and a shifting range for random translation ranging from 1 to 50 units.

By applying these transformations, we successfully generated the requisite input data **(Figure 3 A)**, thereby enabling the subsequent construction of a graph representation for the proteins and ligands, which constituted a fundamental component of our analysis framework.

### 2. Graph Representation for Proteins and Ligands

Following the curation of both the true and random orientations for the proteins and ligands, we proceed to generate graph representations for each molecular entity, as presented in **Figure 2 A and B**. Given that the Graph Neural Network (GNN) exclusively accepts graph inputs, it is imperative that these generated graphs exhibit a high level of feature richness and accuracy. To accomplish this, we leverage the capabilities of PyTorch Geometric and Deep Graph Library (DGL), two powerful frameworks designed for graph handling and feature construction pertaining to proteins and ligands.

**Figure 2:**
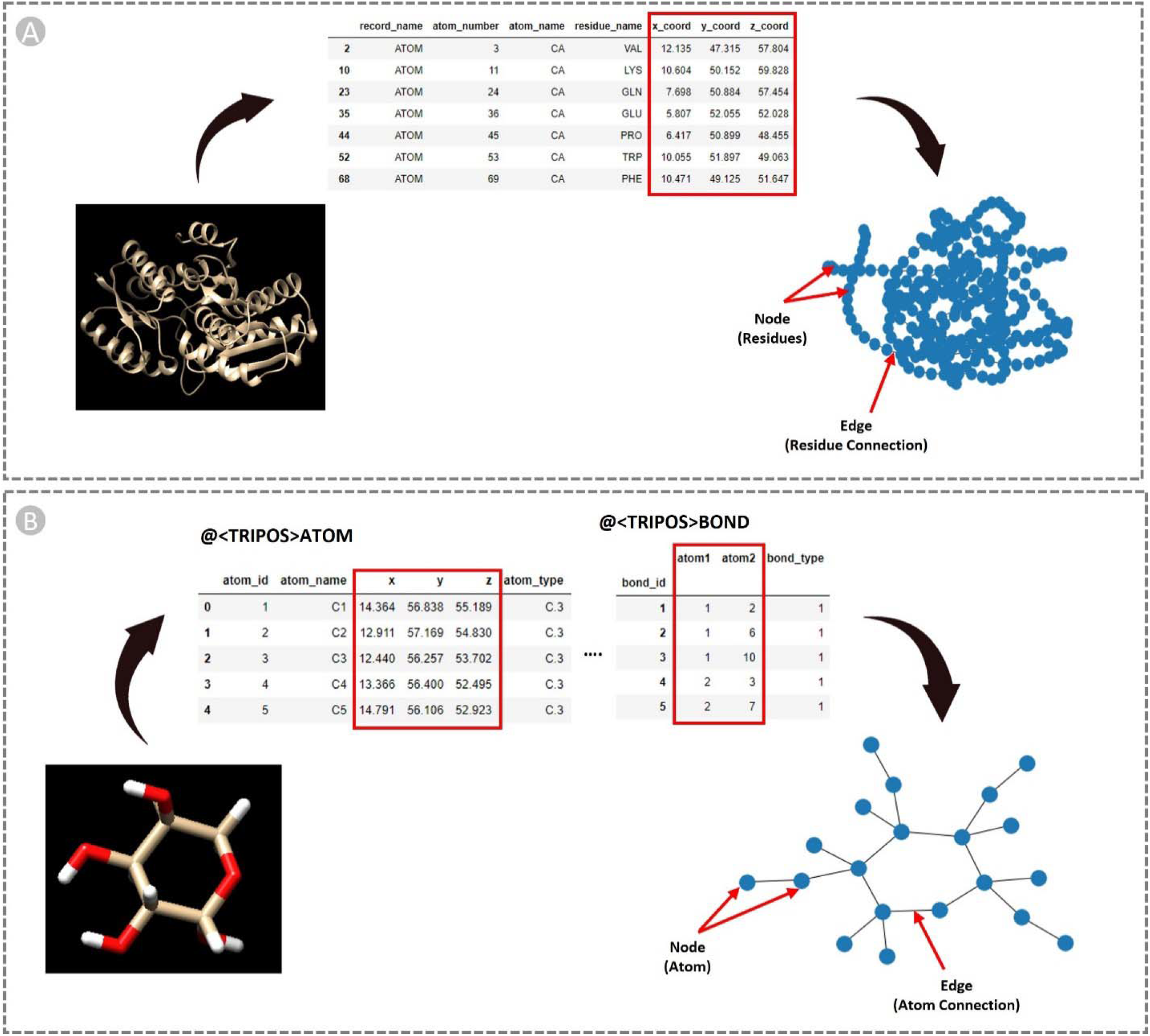
Schematic view of node and edge graph features generation for both receptors (proteins) and ligands.

**Figure 3:**
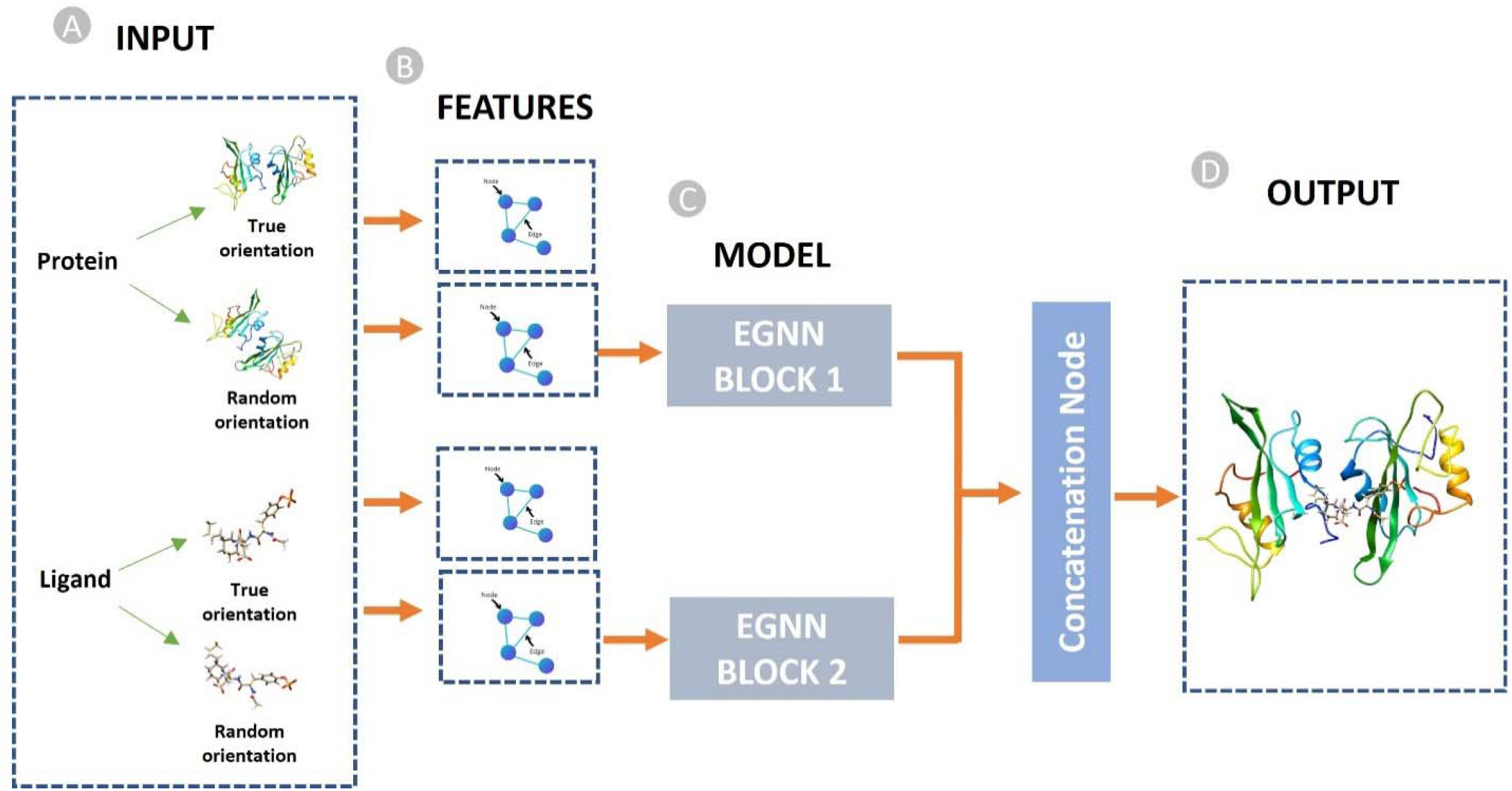
Diagrammatic representation of the overall pipeline: (A) Input data consisting of true protein and ligand structures with random orientations, (B) Generation of graph features based on the input data, (C) Deployment of two Equivariant Graph Neural Networks, and (D) The resultant output depicting the docked protein-ligand complex. The EGNN layers exclusively receive protein and ligands with random orientations, while the true structures are used for the loss function calculation.

In our study, the structural characteristics of the proteins and ligands are portrayed as undirected graphs, denoted as [G_l_ = (V_l_, E_l_)]. In the case of proteins, the nodes within the graph correspond to the residue constituting the molecule, while the edges signify the bonds between these residues (**Figure 2 A)**. While in ligands, the nodes are the atoms of the ligand, and the edges are the connection between these atoms (**Figure 2 B)**. Feature vectors are obtained using RdKit [21] and OGB libraries, which describe the state of a residue and a bond within a molecule. To construct these graphs, we rely on a set of input node and edge features that have been widely adopted in previous research works focused on representing small molecules [1], [22]–[24]. Detailed descriptions of these input features for both proteins and ligands are provided in **Table 1** and **Table 2**, respectively, facilitating a comprehensive overview of their characteristics.

**Table 1:**
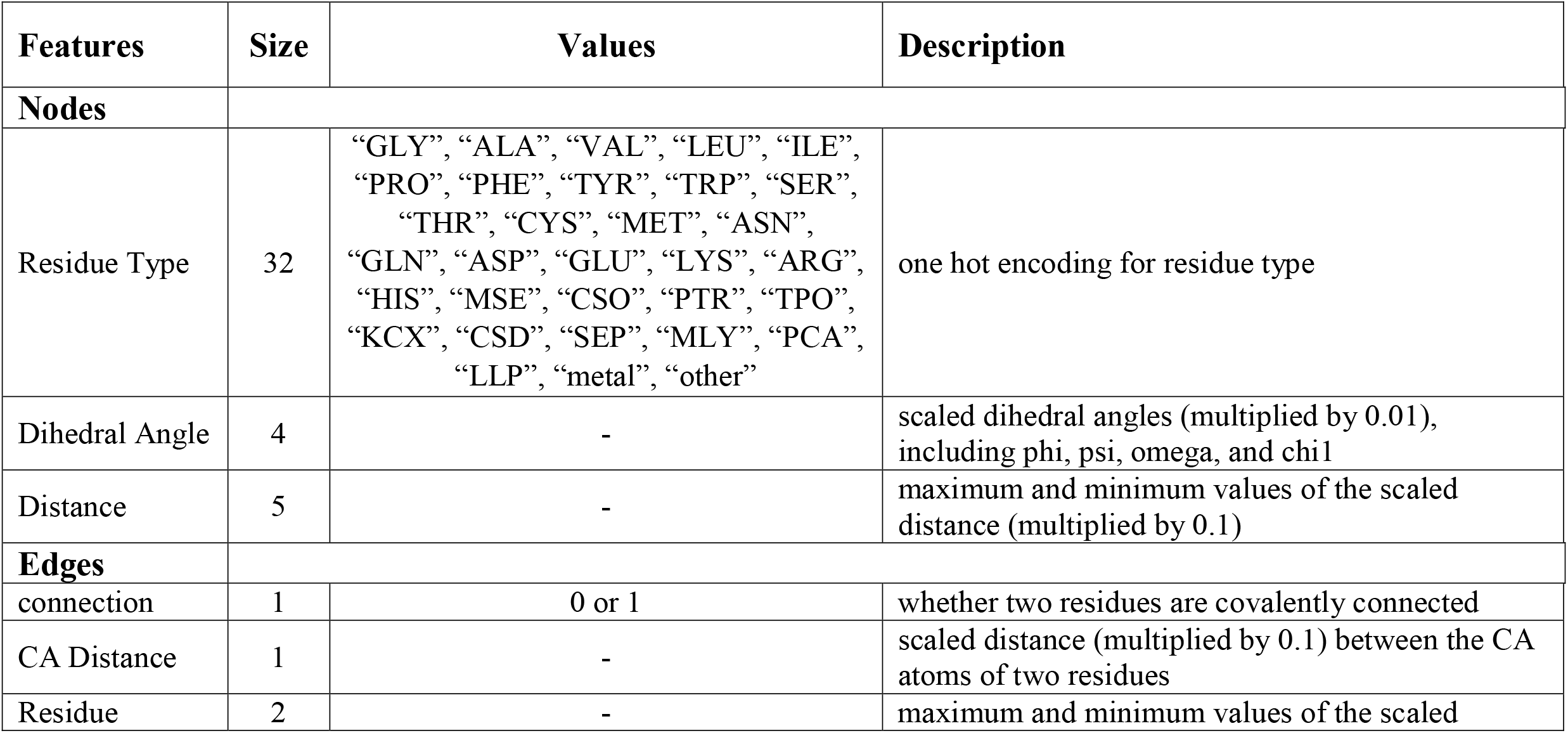
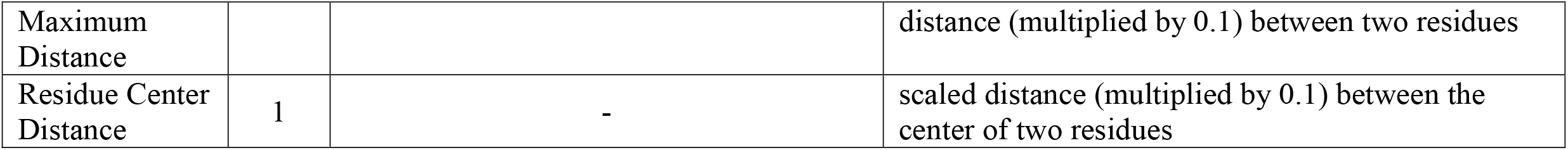
Node and Edge features for protein graphs.

**Table 2:**
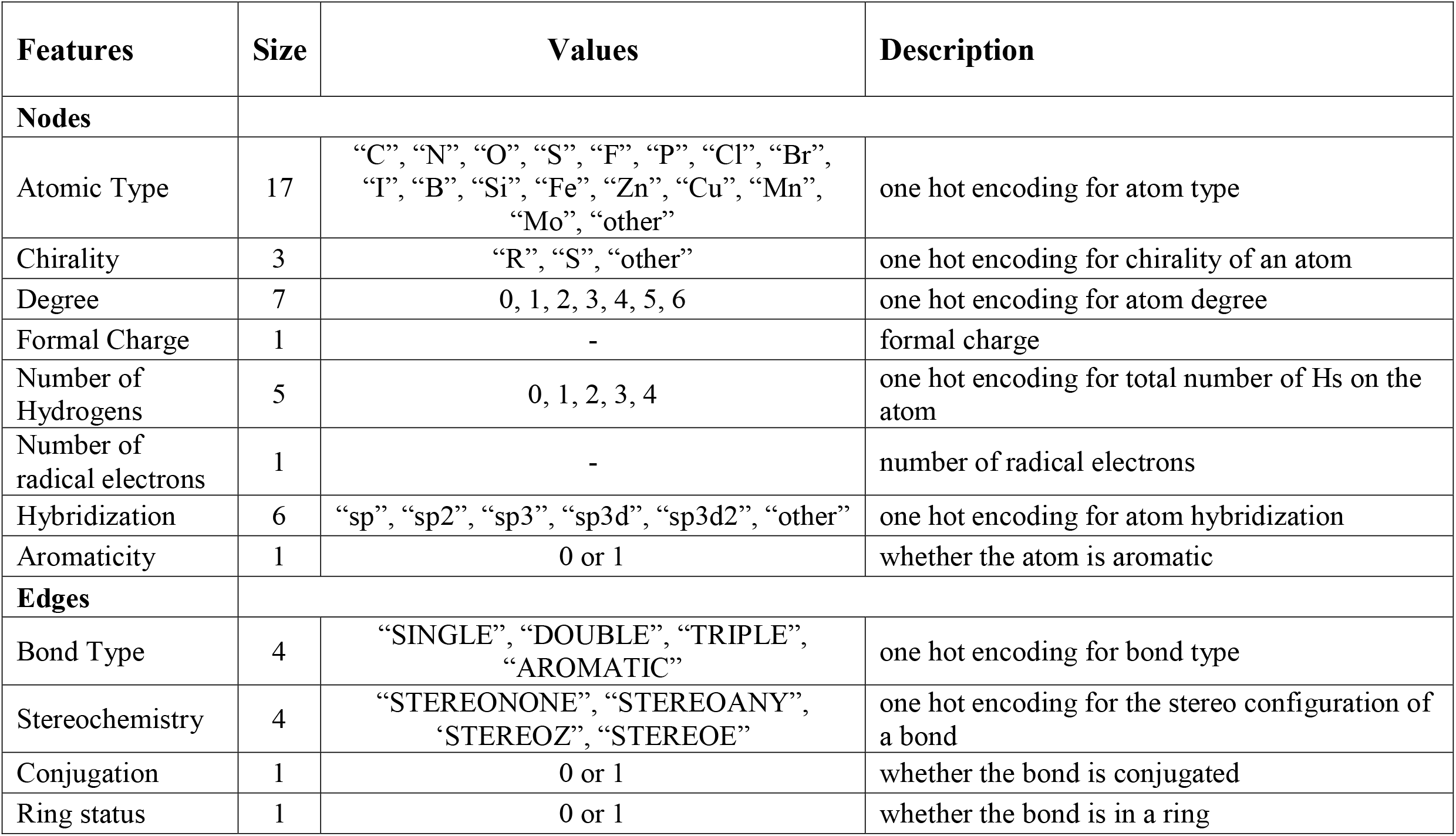
Node and Edge features for ligand graphs.

### 3. Network Architecture Design

We implemented EGNN [10], which possesses the unique ability to maintain equivariance to rotations, translations, reflections, and permutations. Unlike existing graph neural network models that are limited to equivariance in three-dimensional spaces, the EGNN can be easily scaled to higher-dimensional spaces. The overall concept of the pipeline is illustrated in **Figure 3**, demonstrating its efficacy and potential. We harness the power of this model through various experiments conducted in our project.

To train the molecular graph, we employ molecular representations generated by the graph generation step. During each training iteration, the network generates a ligand pose, which is then compared to the original ligand in the protein-ligand complex to calculate the root mean square error (RMSE). To optimize the models, we utilize the Adam optimizer with a batch size of 128, a learning rate of 10^−3^, and a weight decay of 10^−5^.

Two distinct EGNN architectures, namely EGNN1 and EGNN2 **(Figure 3 C)**, are employed to handle individual features of proteins and ligands **(Figure 3 B)**. After predicting the ligand’s orientation, we concatenate the outputs from both models and define the loss function accordingly. By refining the architecture of these models, we ensure that both the protein and ligand are aware of their positions relative to each other. This flexibility enhances the likelihood of achieving improved docked poses which eventually generated the output as the complex of protein-ligand **(Figure 3 D)**.

The model’s design and operation can be comprehended through a series of steps. Initially, the determination of atoms and bonds’ relative positions relies on the fundamental concept of Euclidean distance, providing crucial positional information. Subsequently, message passing occurs across the graphs, facilitating the exchange of information and features among nodes and generating the message M_ij_, which signifies the communication between the i^th^ and J^th^ node.

We consider the graph G = (V, E) with nodes V and edges E, and the feature node embeddings h_i_. A n-dimensional coordinate x_i_ is supposed to be associated with each of the graph nodes. As with GNNs, in this scenario, EGNN will maintain equivariance to permutations on the set of nodes V as well as equivariance to rotations and translations on the set of coordinates x_i_. The network takes the set of node embeddings h^l^ and outputs a transformation on h ^l+1^ and x ^l+1^ as shown in the **Equations (1**-**2)** [10].

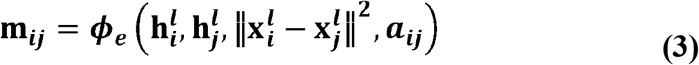

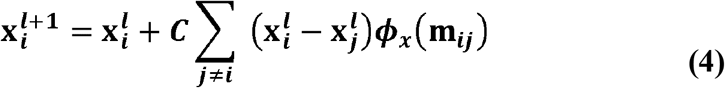

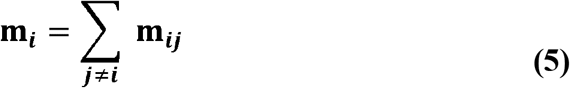

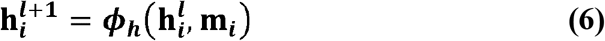

Additionally, there exists the option to compute node attention, allowing for the assessment of nodes and edge features’ significance within the graph. The subsequent phase involves updating the coordinates by passing the received messages through dedicated multilayer perceptions, enabling the inference of atoms’ locations and their desired movements based on the obtained information. Furthermore, capturing the collective moments guides the aggregation of these moments, typically achieved through mean or sum calculations, to serve as the foundation for updating each node. Ultimately, this process leads to the refinement and adjustment of the coordinates, ensuring a comprehensive representation of the molecular system.

In our research, we employ the root mean square deviation (RMSD) as the loss function to assess the plausibility of binding poses. The RMSD between two structures, X_ref_ and X_target_, in the protein-ligand complex is determined by **Equation 7**. The distance, d_ij_, represents the separation between protein atom i and ligand atom j in either X_ref_ or X_target_. The difference between d_ij ref_ and d_ij target_ is summed over all pairs of atoms in a specified contact list, which includes all pairs between heavy atoms of the protein and ligand. The RMSD measurement quantifies the structural changes from the reference to the target structures. By sensitively reflecting the geometrical changes of the ligand relative to the protein, the matrix [d_ij ref_ - d_ij target_] enables the evaluation of binding poses.

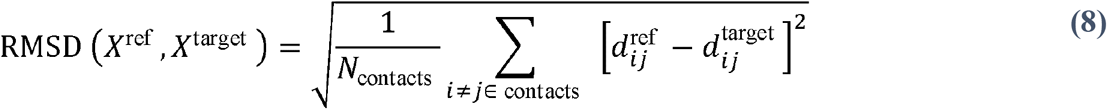

For each protein-ligand complex, we consider the docking pose as the reference structure (X_ref_) and calculate the RMSD of the complex structure at each intermediate predicted pose (X_target_). This approach allows us to assess the quality of the predicted poses and refine our models accordingly.

Through the utilization of the EGNN and the incorporation of the RMSD loss function, our research aims to advance the field of protein-ligand binding pose prediction. By leveraging the unique properties of the EGNN and employing a rigorous evaluation metric, we strive to improve the accuracy and reliability of predicting binding poses. These advancements hold great promise for enhancing structure-based drug design and expediting the discovery of new therapeutic treatments.

## IV. Analysis and Results

The evaluation of the model’s training process was conducted by analyzing loss curves, specifically quantified as the Root Mean Square Deviation (RMSD) between the true and predicted complex structures. Observing the training and validation curves (as shown in **Figure 4 A and B**), it becomes evident that the model exhibits a progressive learning pattern, effectively assimilating information from the input features. This observation serves as a testament to the model’s capability to comprehend and extract meaningful representations from the provided data.

**Figure 4:**
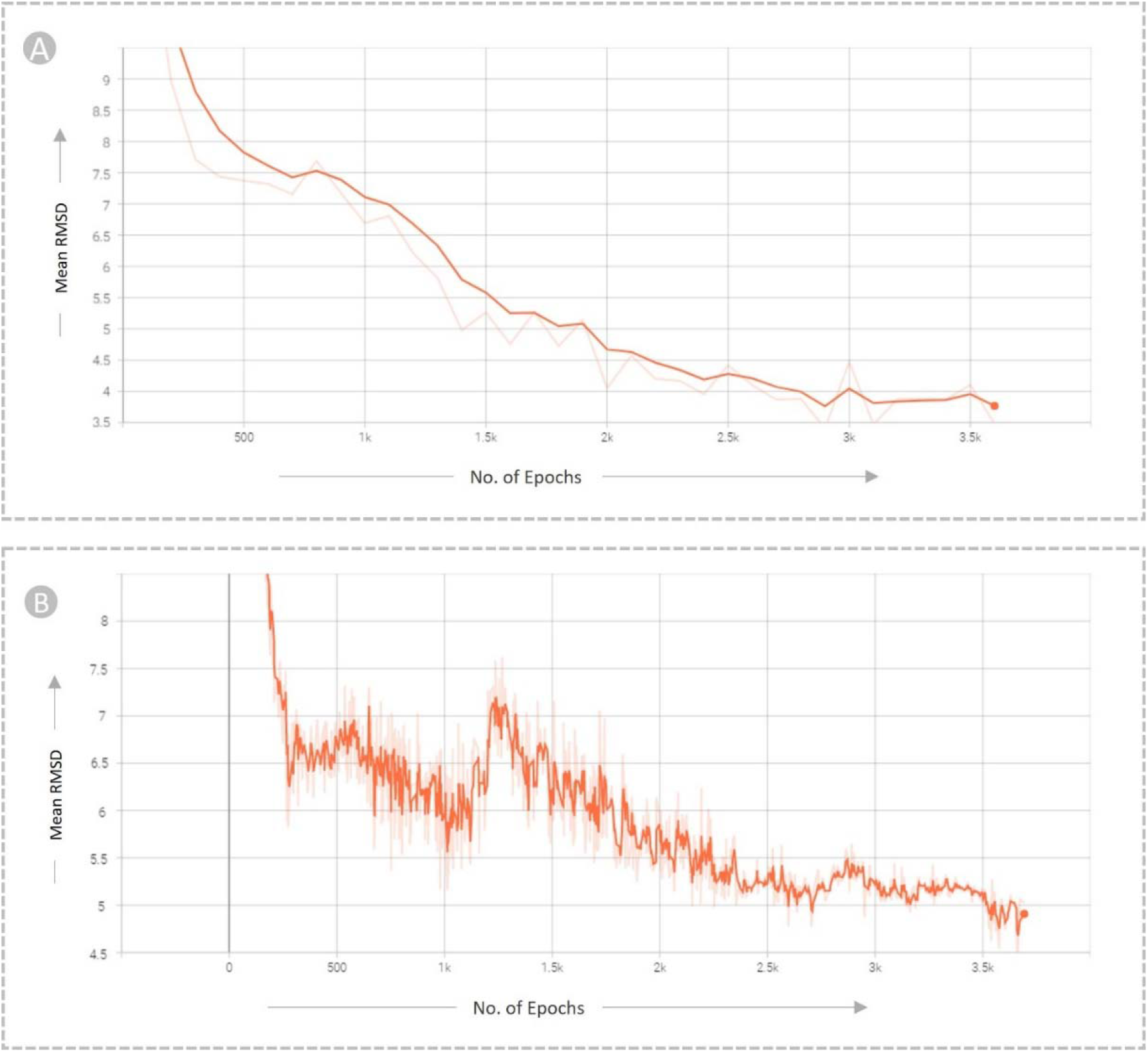
Train Validate loss statistics with Mean RMSD vs number of epochs. (A) Training curve, where the y-axis represents the mean RMSD between predicted and true complex structure, while the x-axis denotes the number of epochs. (B) Validation curve, with the same y-axis and x-axis parameters. Both curves exhibit a decreasing loss trend as the number of epochs progresses.

Following the completion of training, the saved model was loaded to make predictions on previously unseen data from the test set. To provide representative prediction results, we selected two notable examples, namely PDB ID: 1a4m [25] and 1a86 [26]. In these cases, we input the protein structure along with the randomly oriented ligand, allowing the model to predict the x, y, and z coordinates for the docking ligand. Subsequently, we generated two complexes, one based on the true structure and the other utilizing the predicted structure. To comprehensively evaluate the model’s performance, we conducted a visual analysis by superimposing the true structure onto the predicted structure using Chimera’s MatchMaker function [27], [28].

To supplement the visual assessment, we also calculated the TM Score [29] and Root Mean Square Deviation (RMSD) between the two complexes, providing a quantitative understanding of the predictions. The TM score served as a measure of structural resemblance, with a value of 1 indicating a perfect match or identical structures, while a TM score of 0 signifying no structural similarity between the models. In this regard, a higher TM score closer to 1 was indicative of a better performance, highlighting a greater level of structural resemblance between the predicted and true complex structures.

For the specific example depicted in **Figure 5 A** (Case I), the resulting complex comprised 349 residues, exhibiting a remarkable TM score of 0.970 and an RMSD of 2.234 Angstrom. In **Figure 5 B** (Case II), the obtained TM score was 0.909 with an RMSD of 4.509 Angstrom. These quantitative measures further contributed to the comprehensive evaluation of the model’s predictive capabilities, affirming its efficacy in accurately predicting complex structures.

**Figure 5:**
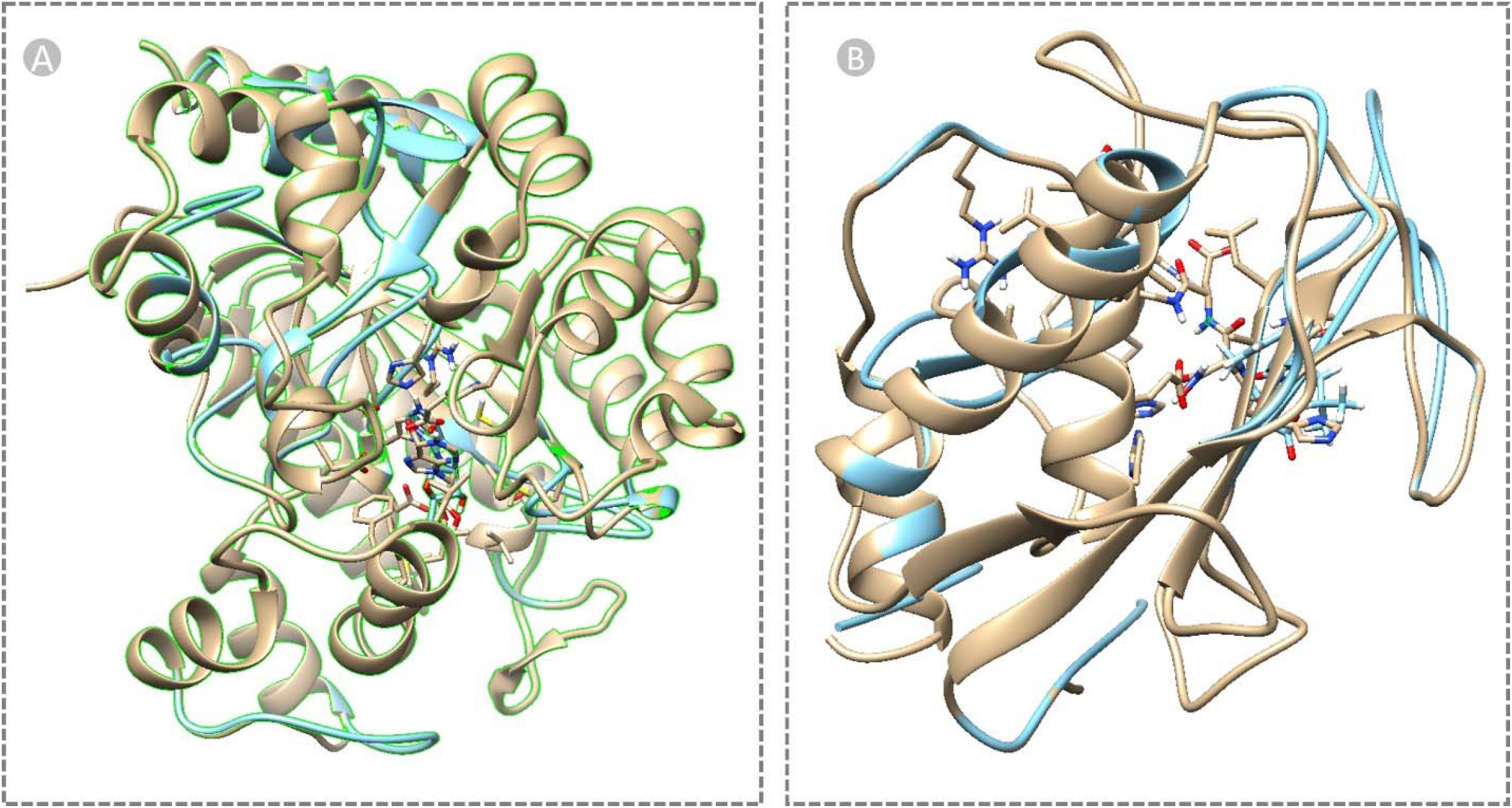
Superimposition of the predicted and actual structures of protein-ligand complexes using Chimera’s MatchMaker function. (A): Predicted (represented in blue) and true (represented in golden) complex structure (PDB ID: 1a4m [25]) with a length of 349 residues. The superimposition analysis yielded a TM score of 0.970 and an RMSD of 2.234. (B): Predicted (blue) and true (golden) complex structure (PDB ID: 1a86 [26]) containing 158 residues. The superimposition analysis for this case resulted in a TM score of 0.909 and an RMSD of 4.509.

## V. Conclusion

The accurate determination of binding poses in protein-ligand interactions holds immense significance in the realm of computational chemistry and in-silico drug discovery. Within the scope of this project, we have introduced an Equivariant Graph Neural Network model to address this problem, which showcases its potential in predicting protein-ligand conformations upon binding. Through rigorous testing, we have demonstrated the efficacy of our approach. Moreover, we envision the future expansion of the prediction framework into a dynamic and versatile architecture, trained on diverse protein-ligand structure datasets, thereby enabling the prediction of optimized binding poses while accurately identifying the binding sites. This progressive trajectory promises to open new avenues and propel the advancement of computational drug discovery, fostering the rapid and efficient development of life-saving therapeutics.

## Notes

### Competing Interest Statement

The authors have declared no competing interest.

